# Grain versus AIN: Common rodent diets differentially affect breeding and metabolic health outcomes in adult C57BL/6j mice

**DOI:** 10.1101/2023.10.16.562483

**Authors:** Lidewij Schipper, Sebastian Tims, Eva Timmer, Julia Lohr, Maryam Rakhshandehroo, Louise Harvey

## Abstract

Semi-synthetic and grain-based diets are common rodent diets for biomedical research. Both diet types are considered nutritionally adequate to support breeding, growth, and long life, yet there are fundamental differences between them that may affect metabolic processes. We have characterized the effects of diet type on breeding outcomes, metabolic phenotype, and microbiota profile in adult mice. Healthy 8-week-old female and male C57BL/6J mice were fed a semi-synthetic or a grain-based diet for 12 weeks and changes in body weight and body composition were monitored. Breeding outcomes were determined. Body fat accumulation of female mice was lower on the semi-synthetic diet than on the grain-based diet. Pregnancy rate and newborn pup survival appeared to be lower in mice exposed to semi-synthetic diet compared to grain-based diet. Both female and male mice showed a profound change in fecal microbiota alpha and beta diversity depending on diet type. Our study shows that type of rodent diet may affect breeding outcomes whilst influencing metabolism and health of female laboratory mice. These factors have the potential to influence other experimental outcomes and the results suggest that semi-synthetic and grain-based diets are not interchangeable in research using rodent models. Careful consideration and increased understanding of the consequences of diet choice would lead to improvements in experimental design and reproducibility of study results.

## Introduction

Preclinical rodent studies are often reported to be subject to poor reproducibility. A lack of appreciation of the extent to which environmental and experimental factors can contribute to rodent health and phenotype may contribute to this issue. The impact of these factors may vary, depending on the parameter and the experimental design, and may contribute to variation both within and between experiments. For example, rodent metabolic health is known to be influenced by a variety of environmental factors including microbiological status (1), social and environmental complexity during and prior to an experiment (2, 3) and previous experiences, such as transportation stress (4). In turn, metabolic health status has the potential to affect a wide variety of other health outcomes in rodents that may be relevant to many types of research, including fertility (5), inflammatory responses (6) and behavioural performance (7). While often overlooked, the type of rodent diet that is used both prior to and during an experiment may be of particular importance.

Semi-synthetic diets, also referred to as purified diets, are produced following standardized recipes, such as those developed by the American Institute of Nutrition (e.g., AIN93-M (8)), using only refined ingredients such as cornstarch, casein, sucrose and soybean oil, with each ingredient generally providing one main nutrient. These diets are often used in nutritional research because they are consistent in production over time. Moreover, the nutrient composition of these diets can be fully customized and controlled, for instance in the design of experimental diets varying in protein or fat source and content and their matched controls (9). However, generic grain-based rodent diets, often referred to as chow diets, are more widely used for studies using rodent models. These diets are produced using ingredients derived from cereal grains (e.g., oats, corn, wheat) and may even contain animal by-products such as fish meal. The ingredients are not refined and each ingredient may contain multiple nutrients. Grain-based diets are subject to variability between production as the levels of nutrients and other non-nutritive compounds, such as phyto-estrogens and pesticides, can vary depending on regional and seasonal influences (9, 10). Both semi-synthetic and grain-based diets are considered to be adequate and meet nutritional requirements to support breeding, growth and a long lifespan for laboratory rodents. While caloric content may be similar, the two diet types are, however, not interchangeable as some of the fundamental differences between the diets are likely to affect metabolic processes and microbiota composition. For instance, the use of purified protein such as casein in semi-synthetic diets may result in higher digestibility and more efficient use of nitrogen compared to grain-based diets with unpurified proteins (11), which may suggest more efficient energy utilization in animals fed semi-synthetic diets compared to grain-based diets. On the other hand, the cereal grains in grain-based diets are rich in soluble fibers that can be fermented by the microbiota in the cecum and colon to produce metabolites, including short chain fatty acids (SCFAs) that can be used as source of energy for the host and are recognized as important modulators of metabolic health (12, 13). In contrast, the source of fiber in semi-synthetic diets is cellulose, which is normally resistant to mammalian enzymes and poorly fermented by the gut bacteria of non-ruminant mammals (14). In recent years, several studies have demonstrated that considerable differences exist in microbiota profile of rodents depending on the types of fiber consumed (15–17).

With the expectation that semi-synthetic and grain-based diets differentially affect energy use and availability, as well as the gut microbiota, these diet types are also likely to differentially affect metabolic phenotypes in laboratory rodents (18). However, systematic and comprehensive experiments exploring the effects of exposure to semi-synthetic diets compared to grain-based diets on metabolic phenotypes in adult mice studies have not yet been conducted. In the current study, we investigated the effects of a semi-synthetic diet versus a grain-based diet on body composition and microbiota composition in adult female and male C57BL/6J mice that were obtained for in-house breeding of offspring intended for other research purposes. Animals arrived in the lab at 8 weeks of age and body weight, body composition and fecal microbiota profile was monitored up to 12 weeks after arrival. The potential effects of diet type on breeding success were also assessed.

## Results

### During acclimatization, body composition and microbiota of female mice are differentially affected based on diet type

Eight-week-old C57BL/6J female mice (S1 Fig), delivered in four separate batches from a commercial animal supplier, were housed socially (two same sex animals per cage) upon arrival and were assigned to a breeding diet that was either grain-based (Grain; Teklad 2916C) or semi-synthetic (Syn; AIN-93G). Body weight and body composition (EchoMRI) was evaluated at arrival and after two weeks, a period representing a typical acclimatization phase. Despite being age-matched there was batch-to-batch variation in body weight and body fat mass upon arrival (absolute body weight and body composition values at arrival and after two weeks are provided in the supplementary data, S1 Table). After two weeks, diet type did not affect the changes in body weight during this period, but Syn diet resulted in a more pronounced loss of body fat mass compared to Grain diet (fat mass (g) change: [F(_1,32_) = 7.14, *p* = 0.01], fat mass (%) change: [F(1,32) = 5.06, *p* = 0.03], Fig 1 A-D). The estimated average daily caloric intake during acclimatization, calculated based on weekly weighings of the food rack per cage, appeared to be increased in Syn compared to Grain fed animals (S2 Table). Microbiota analysis of fecal samples revealed that diet type altered gut microbial composition during the acclimatization phase, as reflected by a decreased alpha diversity (Chao 1 index) in Syn compared to Grain exposed mice after two weeks acclimatization and greater distance between beta diversity clusters of diet groups (S2 Fig). Analysis of changes in microbial relative abundance at genus level indicated a significant interaction between time and diet, indicating that Syn but not Grain diet reduced the relative abundances of Clostridia UCG-014*, Lactobacillus, and* an unknown genus of Muribaculaceae and increased the relative abundance of *Colidextribacter, Faecalibaculum*, an unclassified genus of Lachnospiracea, an uncultured genus of Oscillospiraceae, *Romboutsia and Roseburia.* Grain diet reduced the relative abundance of *Alistipes* and increased *Turicibacter*. The relative abundance of Prevotellaceae UCG-001 *and Ruminococcus* were, however, reduced by both diet types (S2 Fig).

**Figure 1.**
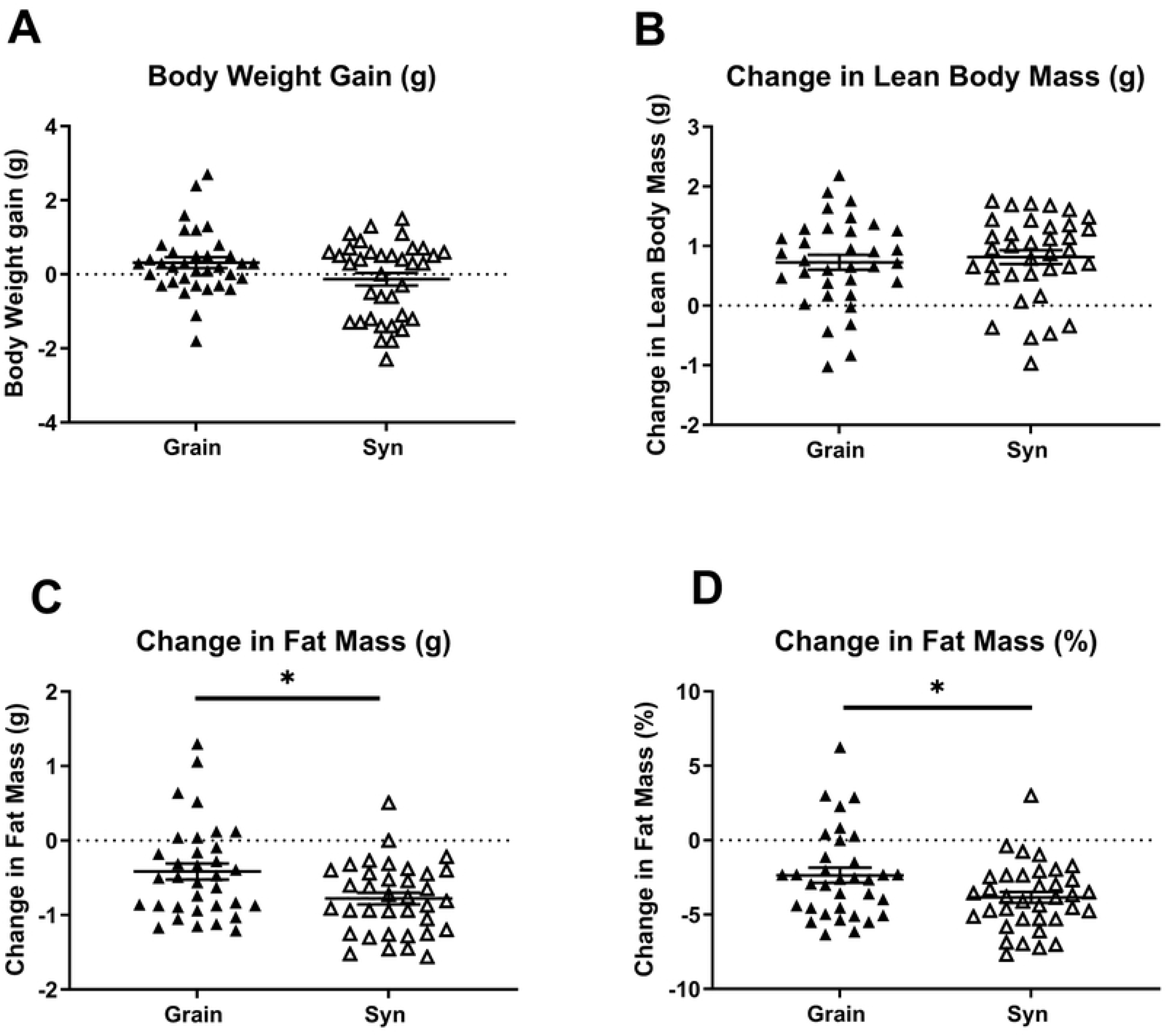
Change in body composition in female mice over 2 week experimental timeline. A) Change in body weight; B) Change in lean body mass; C) Change in fat mass; and D) Change in % mass of female mice during two weeks of exposure to Grain (n = 35) or Syn (n = 36) diet. Data are mean ± SEM. * p < 0.05. Grain: grain-based diet; Syn: semi-synthetic diet.

At arrival, male mice (S1 Fig) were assigned to a standard maintenance diet that was either grain-based (Grain; Teklad 2920) or semi-synthetic (Syn; AIN-93M) and were exposed to the same treatments as females. Males also showed batch-to-batch variation in body weight and body composition upon arrival. In animals that were subjected to Grain some of the batch-to-batch variation could still be detected after the acclimatization phase (S1 Table). While diet type did not appear to affect changes in body weight and body composition and estimated caloric intake of male mice during acclimatization, the changes in microbiota largely paralleled those observed in females in this period (S2 Table and S3 Fig).

### Diet type may affect breeding outcomes

After the acclimatization period, females were mated with males from matching diet groups. Mating resulted in pregnancy in 55.6% of the females on Syn and 75.9% of the females on Grain (*X*^2^ (1, 65) = 2.90, *p* = 0.09, Fig 2A). Within each diet group, the pregnant and non-pregnant females did not show significant differences in body weight or body composition prior to mating (S3 Table). Diet type did not affect litter size at birth (postnatal day 0 [PN0]) (Fig 2B). However, pup survival rate between birth and postnatal day (PN2) was lower in Syn diet litters compared to that of Grain diet litters (84.4% vs 92.8% respectively; *X*^2^ (1, 307) = 5.43, *p* = 0.02, Fig 2C). Litters were weighed for the first time at PN2 and diet type did not affect average body weight of the surviving pups at PN2 (Grain 1.53 ± 0.04 g; Syn 1.50 ± 0.03 g).

**Fig 2.**
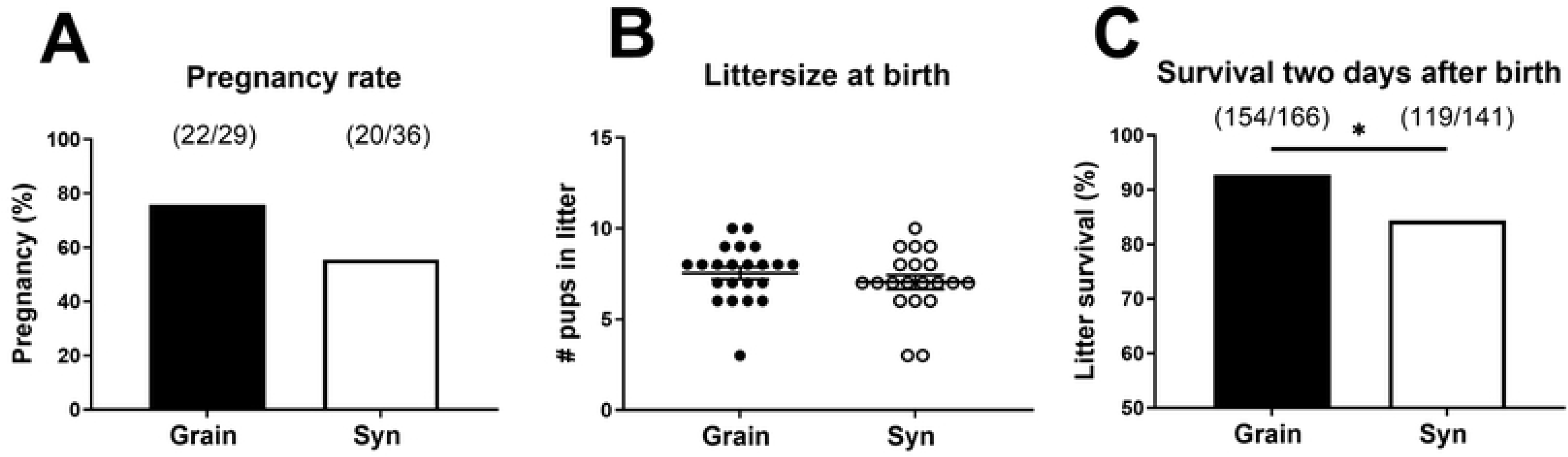
Pregnancy and litter outcomes. A) Pregnancy rate. B) Litter size at birth. C) Pup survival rate two days after birth. Data in B are mean ± SEM. * p < 0.05. Grain: grain-based diet; Syn: semi-synthetic diet.

### Diet type affects body weight and body composition changes over 12 weeks in female mice

Four weeks after arrival, randomly selected (non-pregnant) female mice on the Grain and Syn diet (n = 12/group, representing a group size that is often used for studying metabolic health outcomes in mice) were switched from their breeding diet to maintenance diets (Grain; Teklad 2920, Syn; AIN-93M respectively) and were monitored for 8 weeks. In total, these animals were monitored for a period of 12 weeks after arrival in the lab. Body weight was recorded weekly and body composition was assessed using repeated Echo MRI scans at 4 week intervals (week 0, 4, 8 and 12, Fig 3A). Females showed weight loss or no weight gain during the first 2 weeks after arrival and started to accumulate weight thereafter, with body weight gain of females on Syn remaining lower than that of females on Grain throughout the study (main effect of time [F(_1,21_) = 115.51, *p* < 0.001]; main effect of diet [F(_1,21_) = 6.31, *p* = 0.20], Fig 3B). Lean body mass increased over time and was not affected by the diet type (main effect of time [F(_1,21_) = 29.62, *p* < 0.001]; main effect of diet [F(1,22) = 0.14, *p* = 0.71] (Fig 3C). The animals, however, showed a decline in body fat over the first weeks of the study, and body fat of animals on Syn diet was lower than that of animals on Grain diet (*absolute body fat*: main effect of time [F(_2,22_) = 22.88, *p* < 0.001]; main effect of diet [F(_1,22_) = 29.92, *p* < 0.001]; *body fat (%)*: main effect of time [F(_2,22_) = 18.93, *p* < 0.001]; main effect of diet [F(_1,16_) = 12.76, *p* < 0.01], Fig 3D, E). Body fat appeared to remain stable between 4 and 8 weeks, after which females started to regain body fat. At 12 weeks, only females on Grain diet showed fat mass that was restored to or above baseline values. The estimated average daily caloric intake of females appeared to be higher in Syn compared to Grain fed animals during the first 8 weeks of the study (S2 Table). Animals were sacrificed at 12 weeks of age after overnight fasting, and various organs were dissected and weighed (Table 1). White adipose tisse (WAT) deposition appeared to be reduced in Syn compared to Grain fed animals depending on the adipose tissue depot (total WAT: [F(_1,9_) = 4.54, *p* = 0.06]; *retroperitoneal WAT*: [F(_1,14_) = 7.83, *p* < 0.05]. Relative liver weight was not affected by diet, but adrenal glands of mice were heavier in Syn diet compared to Grain fed females ([F(_1,9_) = 5.29, *p* < 0.05]. Liver triglyceride (TG) content was not significantly affected by diet (Syn 0.19 ± 0.05; Grain 0.15 ± 0.03 TG/BCA protein (mg)).

**Fig 3.**
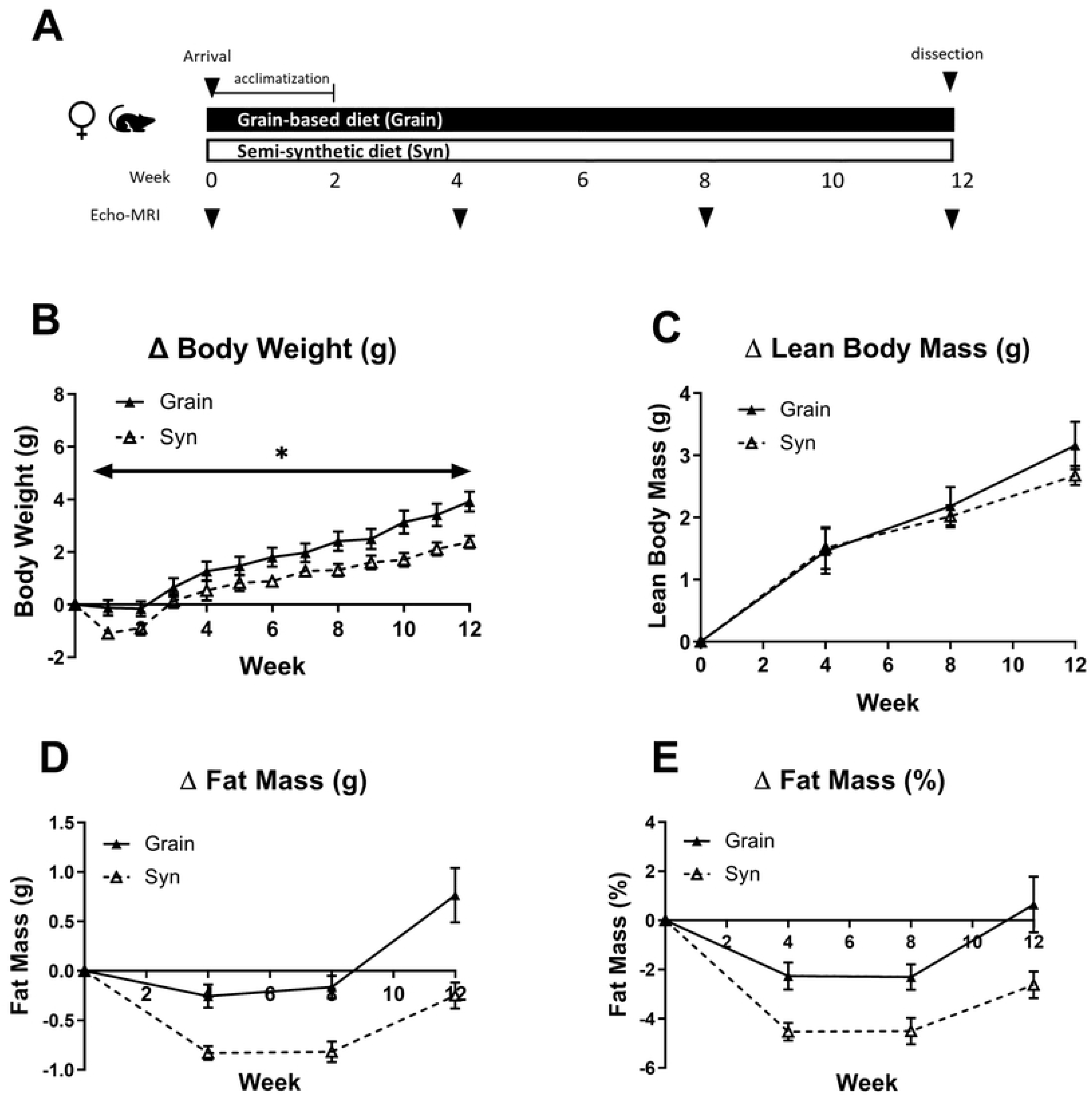
Sampling moments and change in body composition in female mice over 12 week experimental timeline. A) Graphical illustration of timeline illustrating the sampling moments of female mice fed Grain (n = 10 – 12^a^) or Syn (n = 12) diet. B) Percentage body weight gain. C) Change in lean body mass. D) Change in fat mass. D) Change in percentage fat mass. Data are mean ± SEM. * p < 0.05. ^a^ data from 2 animals in the Grain diet group at the 12 week timepoint was missing. Grain: grain-based diet; Syn: semi-synthetic diet.

**Table 1.** Organ weights of female mice after 12 week experimental timeline. Organ weights of female mice exposed to Grain (n = 10) or Syn (n = 12) diet. Data are mean ± SEM. * p < 0.05, ^#^ 0.05 < p < 0.06. BW: body weight; BAT: brown adipose tissue; WAT: white adipose tissue; SCWAT: subcutaneous WAT; ViscWAT: visceral WAT; Syn: semi-synthetic diet. Grain: grain-based diet; Syn: semi-synthetic diet.

|  | Grain | Syn |
| --- | --- | --- |
|  | (n = 10) | (n = 12) |
| **Body weight (g)** | 23.3 ± 0.50 | 23.0 ± 0.33 |
| **BAT (% BW)** | 0.32 ± 0.04 | 0.49 ± 0.07 |
| **Total WAT (% BW)** | 3.33 ± 0.24 | 4.35 ± 0.28<sup>#</sup> |
| **SCWAT (% BW)** | 1.14 ± 0.06 | 1.01 ± 0.06 |
| **ViscWAT (% BW)** | 2.33 ± 0.24 | 1.80 ± 0.17 |
| Gonadal (% BW) | 1.51 ± 0.15 | 1.18 ± 0.17 |
| Retroperitoneal (%<br>BW) | 0.32 ± 0.04 | 0.26 ± 0.02<sup>*</sup> |
| Perirenal (% BW) | 0.50 ± 0.06 | 0.36 ± 0.04 |
| **Liver (% BW)** | 4.46 ± 0.09 | 4.27 ± 0.15 |
| **Adrenals (% BW)** | 0.016 ± 0.001 | 0.020 ± 0.001<sup>*</sup> |
| **Thymus (% BW)** | 0.21 ± 0.01 | 0.24 ± 0.03 |

Male mice on the Grain and Syn diet (Teklad 2920; AIN-93M respectively, n = 12/group) were exposed to the same treatments as females. In contrast to females, there were no significant effects of diet type on body weight gain and body composition development in males over 12 weeks (S4 Fig). At dissection after 12 weeks there were some indications for higher body weight in Syn compared to Grain fed animals but adipose tissue deposition did not appear to be different between groups. Liver weight was reduced, while liver TG content appeared to be higher in Syn compared to Grain fed mice. The estimated average daily caloric intake appeared to be higher in Syn compared to Grain fed animals during the last 4 weeks of the study (S2 Table).

### Diet type affects microbiome and SCFA at 12 weeks in female mice

Cecal short chain fatty acid (SCFA) profile of female mice is described in Table 2. Exposure to Syn diet reduced the weight of the cecal content in female mice ([F(_1,17_) = 10.80, *p* < 0.05]) and also the total SCFA per mL content appeared to be lower in Syn compared to Grain, although this effect did not reach significance ([F(_1,9_) = 4.75, *p* = 0.058]. The relative acetic acid level was increased in Syn compared to Grain fed mice while butyric acid was statistically lower and propionic acid was numerically lower, resulting in higher acetic/propionic and a trend towards higher acetic:butyric acid ratio (acetic acid (%): [F(_1,9_) = 25.189, p < 0.05]; butyric acid (%): [F(_1,9_) = 6.808, p < 0.05]; propionic acid (%): [F(_1,9_) = 4.02, p = 0.08]; acetic:propionic acid ratio: [F(_1,9_) = 8.58, p < 0.05]; acetic:butyric ratio: [F(_1,9_) = 4.83, p = 0.055]).

**Table 2.**
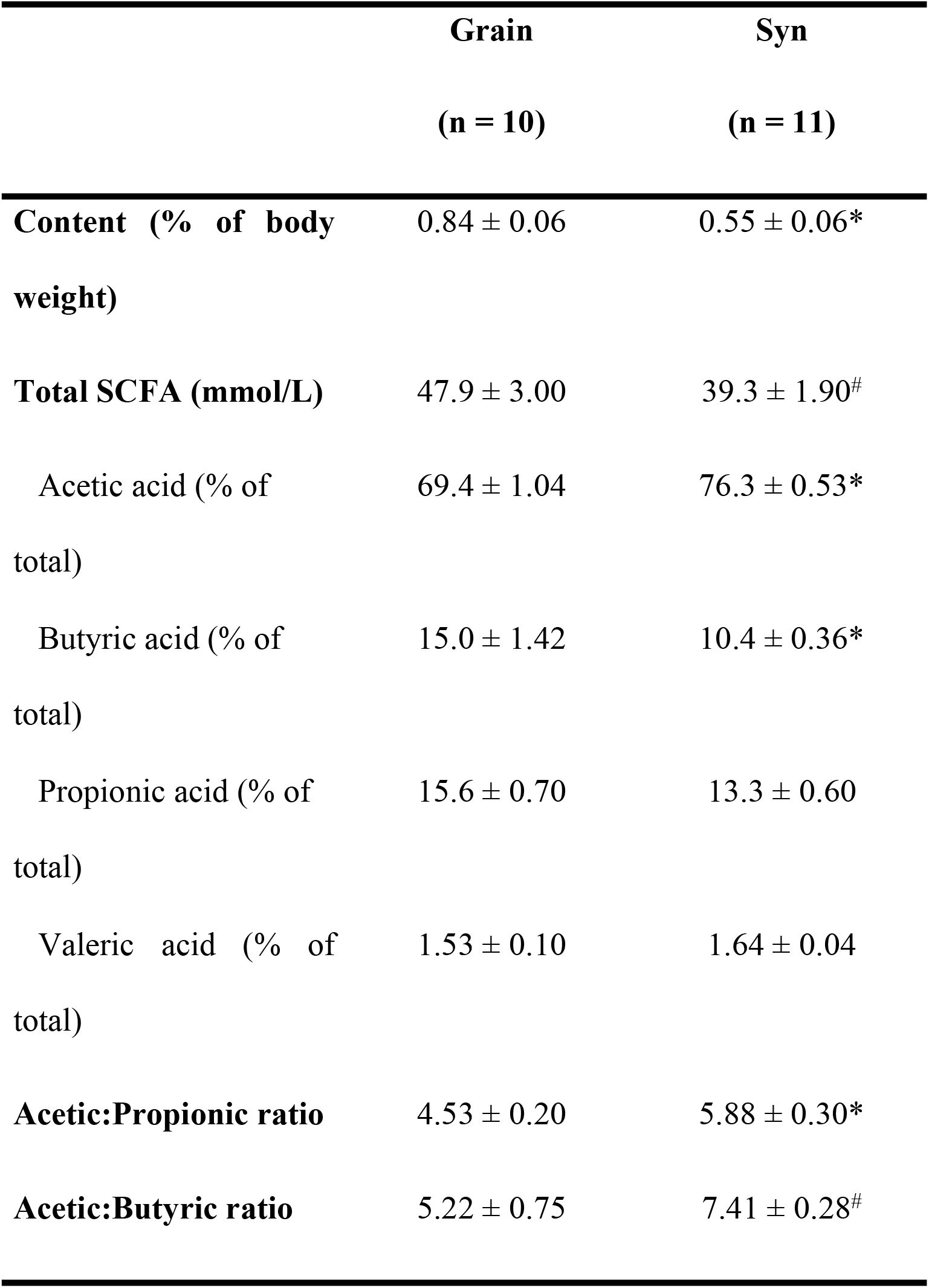
Cecum content and short chain fatty acid (SCFA) profile of female mice after 12 week experimental timeline. Cecum weigh and SCFA profile of female mice exposed to Grain (n = 10) or Syn (n = 11^a^) diet. Data are mean ± SEM.^a^ Cecum of one animal in Syn diet group was not collected during dissection. Grain: grain-based diet; Syn: semi-synthetic diet.

Fecal microbiota analysis in female mice revealed that at 12 weeks, the Syn diet resulted in a signifcantly lower phylogenetic diversity (Chao 1 index, Fig 4A) compared to Grain diet. The PCoA also showed a clear visual separation between the Grain and Syn diet groups at week 12 (Fig 4B). There was a significant diet* time interaction effect observed in the changes in relative abundance of several bacterial genera (S5 Fig). Post hoc testing revealed that while, for many genera, baseline values were not significantly different, the changes over the course of the study were dependent on diet. In both diet groups the genera Rikenellaceae RC9 gut group and *Romboutsia* increased while the genera *Ruminococcus* decreased. But Syn diet lead to significantly higher levels of both Rikenellaceae RC9 gut group and *Romboutsia*, while significantly lower levels of *Ruminococcus* were detected in Syn diet compared to Grain diet. Moreover, exposure to Syn diet resulted in decreased levels of Clostridia UCG-014, *Lactobacillus,* an unknown genus from Muribaculaceae*, Muribaculum* and *Parasutterella* while increased levels of *Colidextribacter*, *Faecalibaterium, Lachnoclostridium,* an unclassified genus of Lachnospiraceae and an uncultured genus from Oscillospiraceae. The reduction in *Faecalibaculum* levels by exposure to Grain diet represents a contrasting shift when compared to the Syn diet. Furthermore, Syn diet seemed to maintain *Alistipes* levels, which were significantly reduced in mice on Grain at 12 weeks. However, the exposure to the synthetic diet did not result in an increase in *Turicibacter* levels, as was observed in mice exposed to Grain diet.

**Fig 4.**
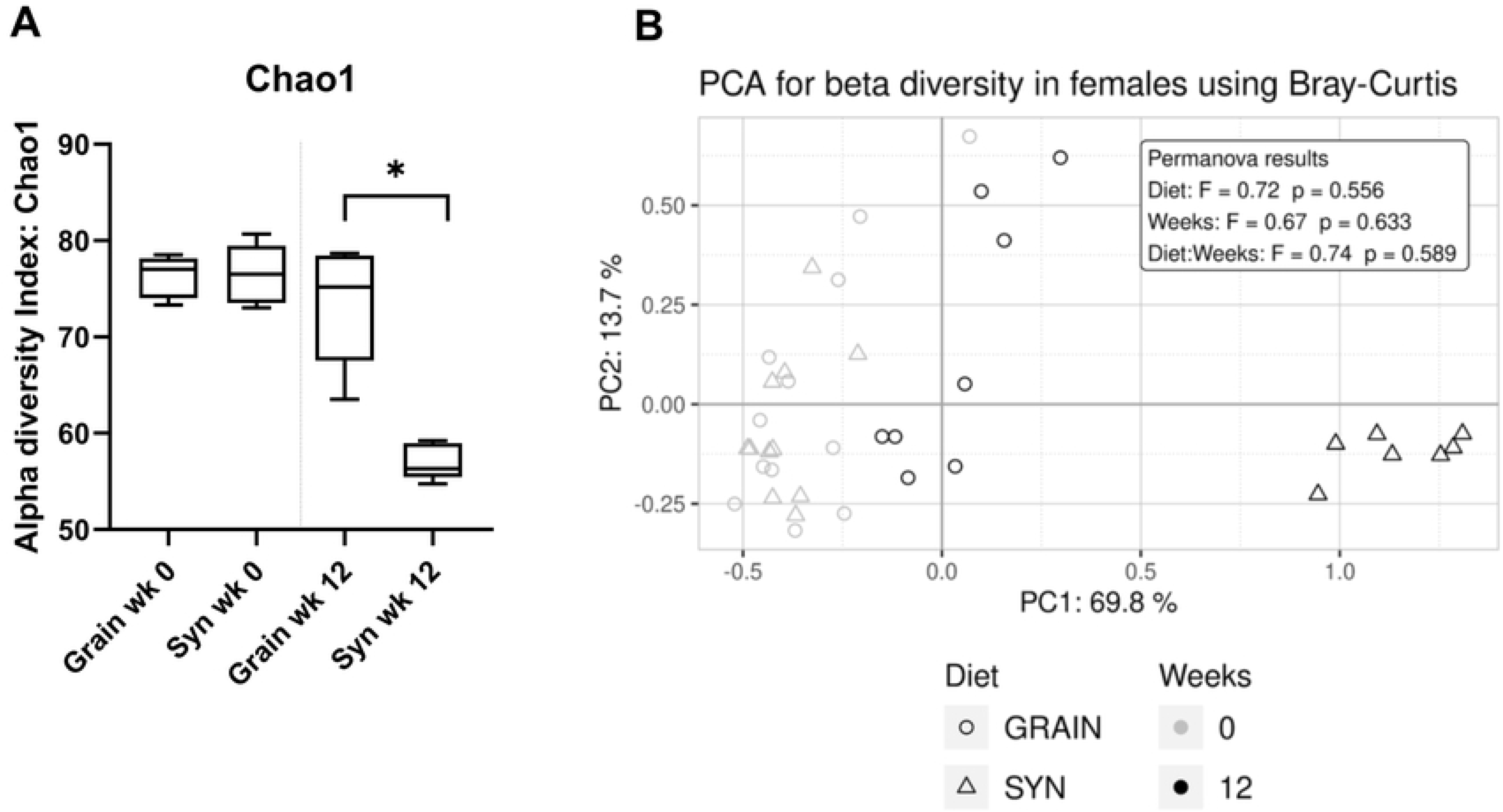
Fecal microbiota composition analyses in female mice at week 0 and week 12. A) Alpha diversity assessed by Chao 1 index of female mice fed Grain (n = 8 – 12^a^) or Syn (n = 7 – 12^a^) diet. B) Beta diversity assessed by principle coordinate analysis (PCoA), using Bray- Curtis distance metrics. Percentage body weight gain. Data are median ± interquartile range. * p < 0.05. ^a^ fecal samples were not collected when mice did not defecate voluntarily at the time of collection. Grain: grain-based diet; Syn: semi-synthetic diet.

In male mice the total SCFA per mL cecum content and the relative butyric acid content was lower in Syn compared to Grain fed mice while the relative acetic acid content and the acetic:butyric acid ratio was increased, which paralelled the effects of diet observed in females (S4 Table). Male mice showed changes in alpha and beta diversity comparable to those observed in females, and most of the changes in the bacterial taxa paralleled those observed in females (S6 Fig).

## Discussion

The current study shows that exposure to a standard semi-synthetic rodent diet compared to grain-based rodent diet impaired breeding outcomes in adult C57BL/6J mice and reduced body weight gain and body fat mass in female mice. These effects were already visible after two weeks of diet exposure and were paralleled by changes in fecal microbiota composition.

While several studies have already compared effects of semi-synthetic and grain-based diet on various health outcomes in rodents, the effects reported on body weight (gain) appear to be mixed. For adult mice, studies report body weight to be higher (19) or lower (11) as a result of semi-synthetic diet compared to grain-based chow feeding, and several studies report no differences depending on diet type (17, 20, 21). This variation in outcomes may be related to differences in experimental design such as chosen mouse strain, the duration of diet exposure and the diet subtypes used. Indeed, different varieties within semi-synthetic and grain-based chows have been shown to influence body weight gain in mice (19). Importantly, the majority of these aforementioned studies were conducted using male mice only, while the results of our study suggest that there are sex differences in the metabolic responses of mice to diet type. In line with our findings, one study reported that semi-synthetic compared to grain-based diet resulted in lower body weight in female mice, while body weight of male mice was not affected (11). Likewise, one study reported that body fat percentage of adult male mice exposed to semi- synthethic diet for 11 weeks was similar to that of mice kept on grain-based chow (22), but no data appears to be available for female mice. While it is often standard practice to monitor body weight in studies as a marker of generic (metabolic) health status, few studies have evaluated effects of diet type on metabolic health status using body composition. The results from our study suggest that, in particular, female mice show changes in body fat percentage in response to diet type.

Exposure to a standard semi-synthetic rodent diet resulted in several changes in fecal microbiota composition compared to exposure to a grain-based diet. Notably, there was a stronger modulation of several bacterial genera in the semi-synthetic rodent diet group, in addition to already obssrvable reductions in diversity during the acclimatization phase. This is in line with a previous study that reported large compositional changes in the microbiota within one week after a switch from grain-based chow to semi-synthetic diet in young adult or mice aged one year (16). Our results show a decrease in alpha diversity in mice on semi-synthetic rodent diet compared to grain-based rodent diet. Other studies have shown associations between lower alpha diversities of the gut microbiota and metabolic disorders, such as obesity, insulin resistance, and metabolic syndrome in both mice and humans (23). This suggest that the changes in microbiota composition observed may be related to changes in energy metabolism and possibly fat accumulation. Several changes observed in the taxa abundances seem to support this. The genus *Alistipes* harbours cultured isolates that are known to be bile-resitant (24) and, therefore, the higher number observed in the Syn group might reflect a change in fat metabolism. *Colidextribacter* was also increased in the Syn group and has been associated with fat accumulation, insulin, and TG in mice challenged with a high fat diet (25). Furthermore, *Romboutsia*, which was also increased in the Syn group, has been associated with lipid profiles and lipogenesis in the liver (26). The Syn mice showed a lowered acetate:propionate ratio, which has been hypothesized to partly explain the impact on energy metabolism differences between mice fed different fibers and propionate, detected in portal vein blood, being able to reach the liver (27). Although here we measured the fecal microbiota and not the cecal microbiota directly, it is reasonable to assume that the cecal microbiota was affected as well, given the observed changes in the cecal SCFA levels. Moreover, a previously reported study showed that significant changes were observed in the cecal microbiota profile of adult male C57BL/6J mice kept on semi-synthetic diet when compared to a grain-based chow for 8 weeks (17). The lower cecum weight in Syn mice observed in our study is in line with a previous study (11), in which it was proposed that the underlying mechanism was linked to a higher osmolaric activity because of the higher fiber content in chow.

### Breeding outcomes

In the current study, using C57BL/6J mice, we observed a reduced percentage of successful pregnancies after timed mating (72 hours) in animals on Syn diet and, while litter size at birth was not different, pups on Syn diet showed reduced survival in the first days after birth. Metabolic health status is one of the factors reported to affect reproductive success and litter outcomes in rodents (5) and we show clearly that this was affected by diet in the current study. However, changes in body weight and body composition in the pre-conception period were not different in females that did not get pregnant compared to those in the same diet group that did get pregnant, suggesting a more direct influence of dietary components on pregnancy outcomes. Protein, energy content and the level of specific components, such as phytoestrogens, within diet type have previously been demonstrated to affect both male and female reproductive function and/or litter outcomes in rodents (28–32). In the current study, care was taken to select diets that were similar in protein and energy content, and phytoestrogen content of the diets was minimal (grain-based) or not present (semi-synthetic diet) to minimize effects of such differences.

However, the SCFA and gut microbiome profiles were strongly affected by diet type and, as these factors have been associated with pregnancy outcomes before (33–35), it is tempting to think that diet induced changes in microbiota may have contributed to the differences observed. More research is needed to confirm this hypothesis; only a limited number of studies have systematically compared breeding outcomes of rodents kept on semi-synthetic diet types to those of rodents kept on grain-based chow, and effects vary between studies. A recent study with Syrian hamsters showed that semi-synthetic diets can strongly reduce reproductive performance (36) whilst a different study, using rats, showed that birth weight was reduced as a result of semi-synthetic diet compared to grain-based diet (37), however, to our knowledge, impaired offspring survival has not been reported. Diet type did not appear to affect fertility, fetal implantation sites and litter size in CD-1 mice that were mated for 10 days (22). While it has been suggested that the effects of diet on breeding outcomes may be species or strain specific (31), other differences in experimental design, such as the duration of dietary exposure prior to mating and the duration of mating, could have contributed to the differences in outcomes between studies. Moreover, pup survival during the first days after birth may be influenced by maternal care and sensitivity to stress (38) that may, in turn, be modified diet type (21). Together, these results warrant further investigation to understand the potential effects of diet type on breeding outcomes in rodents and the underlying mechanisms involved.

### Acclimatization

The results from the current study suggest that in the first two weeks after arrival in a new facility (i.e., acclimatization) the body weight and fat mass of adult female mice were reduced, with stronger effects observed in animals that were allocated to semi-synthetic diet. Animals on semi-synthetic diet required a longer period to regain body weight and fat mass to baseline values. It is important to note that before arrival in the lab, all animals had been kept on a grain-based diet (VRF1, Ssniff) at the commercial animal supplier. While the composition of VRF1 is different to that of the Teklad diet used in the current study, both are grain-based diets and therefore are closer in composition and ingredients compared to a semi-synthethic diet. It is conceivable that a transition from a grain-based to a semi-synthethic diet may have more impact than a transition from a grain-based diet to another grain-based diet; this may have contributed to the differences observed between diet types during and after acclimatization in the current study.

The implementation of an acclimatization phase that allows laboratory animals to adapt to their new environment, including diet, prior to the start of a study is considered common practice. It is typically applied to ensure stabilisation e.g., after transportation stress and increase the likelihood of similar baseline parameters (39). These factors may otherwise have the potential to confound experimental outcomes. For example, in the current study we observed that there was batch-to-batch variation in body weight and body composition in both female and male mice upon arrival despite the fact that animals in the different batches were age matched and were derived from the same breeder facility. These differences were no longer observed in females after the two week acclimatization phase.

The duration of the acclimatization priod that is required varies depending on species, outcome of interest and (previous) environmental factors including diet (40). For laboratory mice, previous studies have suggested between four days and four weeks are required to stabilize to a novel environment; this is based on different physiological stress markers as well as mouse strain (41, 42). Interestingly, some of the batch-to-batch variation in body weight and body composition of male mice at arrival could still be detected after the acclimatization phase in animals exposed to Grain diet. This may suggest that, in addition to diet, the required acclimatization period may also differ depending on sex. At this stage, however, limited information is available on an appropriate length of acclimatization period for C57BL/6J mice in order to preserve and protect breeding and monitoring metabolic health outcomes. More research in this field is warranted.

Attention has recently shifted to the stabilization of microbiota during acclimatization. Relatively rapid changes in the gut microbiota profile of female mice were observed after transport from one facility to another; this appeared to stabilize after approximately one week of grain-based diets and social housing in conventional settings (39). Another study showed that the transition from a grain-based chow to semi-synthetic diet was also able to signifcantly change microbiota profile of mice within one week (16). In our study, animals were both transported to a new facility and switched to a different diet type and, in line with the aforementioned studies, we observed that at least 10 genera were affected after two weeks of acclimatization to the semi-synthetic diet, and to lesser extent to the grain-based diet, in female mice. However, interestingly, we observed a few additional bacterial genera were affected in the Syn groups after 12 weeks, suggesting that the microbiota profile continues to develop, even after 2 weeks of acclimatization in IVC conditions, and that the effects are different per diet type. Taken together, the results from the current study suggest that two weeks of acclimatization on either diet type may not be sufficient to stabilize body weight, body composition and microbiota profile in adult C57BL/6J mice that are used for breeding and/or studies focusing on metabolic health. While full stabilization and standardization of all factors would be challenging, closer monitoring of the on-going changes in microbiome profile and body composition during the acclimatization phase in specific models and facilities may help explain variation between study outcomes and improve future study design.

### Limitations

Data on caloric intake per animal was estimated rather than precisely measured and should be interpreted with care. Caloric intake could not be determined at the individual level as animals were housed in pairs. Moreover, spillage of food from the cage rack could not be prevented. While there are more reliable methods for the assessment of individual caloric intake, including individual housing in cages with food hoppers equipped for food intake registration, these methods were not used in the current study. Instead we prioritised social housing over individual housing, as social isolation may induce stress in mice and energy intake during social isolation may not be representative of energy intake during social housing (3). Interestingly, several studies report a lower intake of energy per g body weight in semi-synthetic compared grain-based chow fed rodents while body weight remains unaffected or is higher (17, 19, 21, 43, 44). This may suggest more efficient utilization of energy from semi-synthetic diets. In contrast to these findings, however, the results of the current study suggest that, if any, estimated caloric intake is higher when animals are exposed to Syn compared to Grain diets. More specialized research that includes careful monitoring of energy intake and energy expenditure per individual would be needed to confirm these observations.

In the current study we explored the gut microbiome because diet induced changes in microbiota and SCFA profile may act as potential modulators of metabolic health. However, recent work suggests that in addition to gut bacteria, viruses and fungal communities of molds and yeasts in the gut, collectively referred to as the mycobiome, can be shaped by dietary components and contribute to metabolic phenotype of laboratory mice (45). It remains to be confirmed in future studies whether the gut mycobiome is affected by semi-synthetic vs grain- based diet feeding.

## Conclusion

Semi-synthetic and grain-based diets can differentially modulate various health outcomes in laboratory mice including metabolic phenotype and breeding outcomes. Effects on metabolic phenotype appear to be sex dependent, with semi-synthetic diet compared to grain-based chow leading to lower body weight and reduced adipose tissue deposition in female mice already after two weeks of acclimatization. Exposure to the different diet types results in rapid changes to the fecal microbiota profile. The results of the current study contribute to the ongoing debate about grain-based and semi-synthetic diet types that has ensued following an increase in awareness about the potential (confounding) effects that the use of different diet types can have on study outcomes (9, 18, 46–48). Careful consideration of the potential influence of rodent diet type on metabolic phenotype and other health outcomes may help optimize future experimental design and data collection in a wide range of preclinical studies, not limited to nutritional intervention experiments. In addition, more information about diet induced differences in phenotype may help explain some of the variation in outcomes between different studies that use different diet types.

## Materials and Methods

### Ethics statement

This study was conducted under an ethical licence of the national competent authority (CCD, Centrale Commissie Dierproeven) including a positive advice from an external, independent animal ethics committee (DEC consult, Soest, The Netherlands), and all animal procedure were prospectively approved by the Animal Welfare Body, following principles of good laboratory animal care, securing full compliance to the European Directive 2010/63/EU for the use of animals for scientific purposes (protocol approval code: INUTEX-19-11 02_001_GNL_LS). The C57BL/6J mouse strain was selected for this study since this strain is one of the most common inbred mouse strains used in biomedical research.

### Animals and care

In total, 32 male and 72 female C57BL/6J mice (SPF, 8-week-old), were obtained from Charles River Laboratories, Saint-Germain-Nuelles, France). Animals arrived in the lab in four batches separated by 4, 5 and 12 weeks respectively. Animals were marked for identification purposes (ear clip) upon arrival and were housed in same sex pairs throughout the study unless specified otherwise. Females were randomly assigned to pairs after arrival. We deliberately obtained male animals in pairs that were littermates and that had been housed together from weaning onwards in an attempt to minimize aggression between adult male cage mates. The males were kept in the same pair after arrival until breeding (see below) and were reunited with their former cage mate after breeding. Males remained socially housed with their respective male cage mate throughout the rest of the study, and fighting incidence was closely monitored. All animals were kept in IVC polycarbonate type II cages randomly positioned on two racks in the same room (21 ± 1°C; 50 ± 5% humidity; 12/12 light/dark cycle with lights on at 07:00). All procedures took place during the light phase. The cages were equipped with bedding (Aspen wood shaving), two plastic transparent tunnels (red and yellow) and atransparent plastic igloo (Datesand, Manchester, UK) as cage enrichment items and aspen wood wool as nesting material. Animals were transferred every other week to a fresh cage with bedding; the cage enrichment items and wood wool were not refreshed and were transferred from the old to the new cage. Cages were opened one at a time in a flow cabinet and working surfaces and handler gloves cleaned with 70% Ethanol between opening individual cages. Male cages were always handled before females cages, and garments were refreshed in between handling sexes to minimize risk of transfer of female scent to males. Additionally, the order of handling cages was randomized using a random sequence generator app. Food (see below) and sterile water were available ad libitum unless specified otherwise. Animals were picked up using tubes rather than by the base of tail to reduce stress (49).

### Dietary exposures

All animals had been kept on a standard grain-based diet VRF1 (Ssniff) at Charles River, the commercial animal supplier. Upon arrival animals were randomly assigned to either semi-synthetic diet (Syn; AIN-93G for females, n = 36 and AIN93-M for males, n = 16 (8)) or grain-based chow (Grain; Teklad 2916C for females, n = 36 and Teklad 2920 for males, n = 16). After acclimatization and timed mating (see below), non-pregnant females (12 per dietgroup) were switched to the respective maintenance diets (AIN93-M and Teklad2920) and were kept in the study for 8 more weeks (Grain, n = 12; Syn n = 12). Similarly, males (12 per diet group) were also kept in the study after mating. Due to the difference in colour and texture of grain-based chow and semi-synthetic diet the researchers could not be blinded to exposure to diet type upon collection of *in vivo* parameters. *Ex vivo* analyses and data processing was performed by researchers unaware of group allocation.

### Breeding procedure and outcomes

After two weeks acclimatization in the lab, females were time-mated. Cages were not cleaned in the week prior to and after breeding to reduce stress. Three days before mating a handful of soiled bedding from a male cage was placed in a female cage with matching diet type. After three days males were removed from their home cage and placed in a cage with two females (one male per cage). After 72 hours, the males were returned to their (soiled) home cage in pairs and were closely monitored for signs of fighting. Animals were used only once for breeding. Pregnant females were housed individually from 17 days after breeding and non-pregnant females were re-assigned to another non-pregnant cage mate from the same batch and diet group. Birth of litters was monitored every morning between 8:00 am and 10:00 am and the number of pups in the litters at birth was recorded. At PN2, the number of pups in the litter was recorded again. After PN2, the dams and litters were used for other research and were no longer monitored as part of the current study.

### Body weight, body composition and caloric intake

Body weight was recorded every other week for all animals. Body composition of the mice was analyzed non-invasively by magnetic resonance imaging using an EchoMRI-100™ analyzer (EchoMRI Medical Systems, Houston, TX), assessing lean body mass (g) and fat mass (g) of the animal. Relative fat mass and relative lean body mass were calculated based on body weight (g). EchoMRI scans were taken at week 0 (arrival), 2, 4, 8 and 12. Mice were guided into a 50 cm long EchoMRI tube with an inside diameter of 28 mm and a smaller tube was inserted after to prevent the animal from escaping from the tube. The tube was inserted in the echo apparatus and a scan was made within 2-3 minutes. Animals were returned to their home cage after the scan. Caloric intake per animal per day was estimated by weighing of the food on the cage rack (at least once per week upon supply of fresh food) and dividing the total calories consumed per cage by the number of animals in the cage and the number of days that had passed since the last weighing. The average daily consumption was calculated for the acclimatization period (average of 2 weeks), and for each 4 week interval in the total 12 week follow up period.

### Fecal sample collection

Fresh fecal samples were collected at week 0 (arrival), 2 and 12 and after fasting, just prior to dissection. Mice were placed individually, for a maximum of 15 min, in an empty type II cage without bedding until they produced a fecal sample. Samples were collected, snap frozen and stored at -80°C until further processing. No samples were collected in situations where the mouse did not produce a fecal sample within 15 min.

### Tissue collection and plasma measurements

After 12 weeks of diet exposure, animals were provided with limited amount of food at 16:00 (2.5 g/animal) and were deeply anaesthetized the next morning using isoflurane inhalation followed by heart puncture or orbital bleeding and decapitation. Blood glucose was determined using a commercial blood glucose meter and test strips (Accu-Chek Performa, Roche Diabetes Care, Inc) and after centrifugation plasma was snap frozen and stored at -80°C until further processing. Adipose tissue depots (interscapular brown; subcutaneous, gonadal, perirenal, retroperitoneal white), and organs (liver, thymus, adrenal glands, cecum) were dissected and weighed.

### SCFA analyses

Cecum samples were diluted 1:10 according to weight in pre-cooled phosphate-buffered saline (PBS). Samples were vortexed 3 times for 30 sec and centrifuged at 4°C for 5 min at 15 000 relative centrifugal force (RCF). The supernatant was collected and 200 µL was used for SCFA analysis. The following SCFAs were quantified on a Shimadzu-GC2025 gas chromatograph with a flame ionization detector and hydrogen as mobile phase: acetic, propionic, n-butyric, iso-butyric, n-valeric, and isovaleric acids. Quantification was performed by using 2- ethylbutyric acid as an internal standard and generating a calibration curve from the peak area after which the concentration in the samples was calculated.

### Liver triglycerides

Liver triglycerides were determined in liver homogenates prepared in buffer containing 250 mM sucrose, 1 mM EDTA, and 10 mM Tris-HCl at pH 7.5 using a commercially available kit (Instruchemie, Delfzijl, The Netherlands) according to the manufacturer’s instructions.

### Fecal DNA extraction

DNA extraction was performed from 0.1 g of fecal sample using QIAamp PowerFecal Pro DNA Kit (Qiagen, Hilden, Germany). The solid fecal sample was added into PowerFecal Pro tubes containing pre-filled beads and the extraction was performed according to the manufacturer’s instruction with additional bead-beating step (ensruing a total of three bead-beating steps) and RNase A treatment. After the last bead-beating step the fecal supernatant was centrifuged at speed of 15,000 x g for 1 min, followed by addition of 8 µL of DNase-free RNase (100 mg/mL) and incubated at room temperature for 2 min, after which the manufacturer’s instructions were followed untill the end of the procedure. 200 µL of CD2 solution was added to the fecal supernatant, vortex-mixed for 5 sec and centrifuged at speed of 15,000 x g for 1 min. The fecal supernatant was then transferred to a clean 2 mL microcentrifuge tube, added with 600 µL of CD3 solution and vortex-mixed for 5 sec. The lysate was then loaded into the provided MB Spin Column. The DNA attached to the silica membrane in the MB Spin Column was washed in two centrifugation steps by different wash buffers i.e. solution EA and solution C5. Finally, the purified DNA was eluted in 50 µL of elution buffer (Solution C6). DNA quantity and quality (A260/A280 and A260/A230) were measured using a NanoDrop 2000 spectrophotometer (Thermo Fisher Scientific, USA) and the remaining DNA was stored at -80°C until use.

### Sequencing

The V3-V4 region of the 16S rRNA gene was PCR-amplified, with primers Bact-0341F (5′- CCTACGGGNGGCWGCAG-3′) and Bact-0785R (5′-GACTACHVGGGTATCTAATCC-3’) (50), and sequenced on the MiSeq platform (Illumina) as described previously (51) using the 2 x 300 bp paired-end protocol. Sequencing was performed by Nutricia Danone in Singapore. In each sequencing run 5% of PhiX (Illumina) was included as an internal control. The read pairs were demultiplexed and trimmed (q>20) before being merged using Paired-End reAd meRger (PEAR) (52) v0.9.2. Merged reads with q>25 over a window of 15 bases, no ambiguous bases and a minimal length of 300 were retained. These were dereplicated and counted using mothur v.1.41.1 (53) and reads with a low abundance (less than 2 reads over all samples) were discarded. Chimeras were removed using VSEARCH v.2.3.4 (54), using the ChimeraSlayer reference database (55) as reference. Reads which contained PhiX or adapters as defined in Deblur v.1.1.0 (56) (part of QIIME2 (57)) were eliminated. Taxonomic assignment was performed using the RDP classifier v.2.2 (58) against the SILVA v. 138 (59) database. Reads with eukaryotic or chloroplast assignments, as well as reads with a low relative abundance up to 0.0005% in all samples were excluded from further downstream analysis.

### Sample size and statistics

The sample size that was used for each phase of the current study is illustrated in the supplementary materials (**Supp Materials 1A, B**). The animals that were used in the current study were originally obtained as breeder animals to generate offspring intended for other research purposes. Sample size calculations were not performed. There were 72 females (36 Grain; 36 Syn) and 32 males (14 Grain; 18 Syn) used for the data collected during the acclimatization and breeding phase of the study. For the 12 week follow-up phase, 12 animals per group were used. This number was based on the smallest number of male and female mice for each of the diet group that were still available after breeding and that were eligible for long term follow up (i.e., socially housed and non-pregnant).

There were three dropouts in the week after arrival: one female and one male with malocclusion were taken out of the study upon arrival and one male was found dead in the home cage after one week, the cause of death remains unknown. This resulted in two males being housed individually prior to breeding rather than in pairs and one occasion where three females were housed together in one cage prior to and during breeding. As individual compared to social housing is known to affect body composition in mice, data from these animals were excluded from analysis (3). One pair of males housed socially after breeding were removed from the study due to excessive fighting (S1 File). After nine weeks one female on Grain diet presented with a skin lesion due to excessive grooming and had to be euthanized. To prevent individual housing the cage mate was also excluded from the study.

All data were analyzed for males and females separately since body weight and bodycomposition are strongly dependent on sex. For body weight, body composition, organ parameters and breeding outcomes all statistical analyses were performed using IBM SPSS Statistics 19. Effects of diet type on dichotomous variables were analyzed using Chi-square test. Other data were analyzed using linear mixed models: Effects of diet type on changed body weight and body composition at 2 weeks (acclimatization) and on organ and plasma parameters at 12 weeks were analyzed using diet as fixed factor, excluding missing datapoints (e.g. dropout or lost samples). Effects of diet type on changes in body weight and body composition over 12 weeks were analyzed using diet, time and the interaction between these variables as fixed factors, excluding data at missing timepoints (dropout). Individual animals were considered as statistical units, however, as the study included multiple batches of mice and mice were always housed with two animals per cage throughout the study, analyses were run with cage as random factor where appropriate. Significance was set at p < 0.05. Statistical trends were reported in case of a p-value between 0.05 and 0.06. Data are presented as mean ± SEM unless otherwise specified.

For analysis of fecal microbiota, rarefaction was applied to the taxa by phyloseq (60) and vegan packages (61) in R v4.0.2 (A language and environment for statistical computing https://www.R-project.org/ (2018)) in order to perform alpha diversity calculations using the Chao1 and Shannon index metrics. The beta diversity was computed over all samples using the vegan R v4.0.2 package. Statistical significance of differences in alpha diversity were assessed with pairwise_wilcox_test function from the rstatix package in R v4.0.2 (62) followed by Benjamini-Hochberg p-value adjustment per timepoint. Statistical significance of differences in beta diversity were assessed using the permutation ANOVA function adonis2 from the package vegan in R. Genera with a minimum mean relative abundance of 0.5% across all samples were tested for differential abundance between diets. Differential abundance was performed with generalised linear models with mixed effects on the sequencing counts using the glmmTMB package v 1.1.2.3 in R v4.0.2 (63) followed by Anova.glmmTMB applying the Chi Squared test for significant differences. The resulting p-values were corrected using Benjamini-Hochberg. Models containing significant interaction effects were followed up with a posthoc using the emmeans package v 1.7.5 (64) in R. A correction for the variables Cage and Cohort were only applied in each genus’ model when they showed differential abundance as main effects in univariate models to reduce the risk of overparameterisation. After adjustment, a p-value < 0.05 was considered significant for all statistical tests applied to the sequencing data.

## Acknowledgments

The authors would like to acknowledge the many people who contributed to this study either in the design phase, the execution or in the analysis: Cleo Arkenaar, Martin Balvers, Eline van der Beek, Martijn Breeuwsma, Nicole Buurman, Francina Dijk, Miriam van Dijk, Jessica Freesse, Johanneke van der Harst, Andrea Kodde, Stephan Pouw, Hanil Quirindongo, Noela Schaap, Heleen de Weerd, and Tjalling Wehkamp.

## Supporting information

**S1 Fig. Flow diagram of study.** A) Flow diagram of study for female mice. B) Flow diagram of study for male mice. Grain: grain-based diet; Syn: semi-synthetic diet.

**S2 Fig. Fecal microbiota composition analyses in female mice at week 0 and week 2.** A) Alpha-diversity of female mice fed Grain (n = 10 – 13^a^) or Syn (n = 8 – 14^a^) diet assessed by Chao1 index. B) Beta-diversity assessed by principle coordinate analysis (PCoA), using Bray-Curtis distance metrics. C-O) Box plots of bacterial taxa (at genus level) at week 0 and week 2 with significant interaction, as assessed with generalized linear models with mixed effects on the sequencing counts followed by Chi Squared test. The resulting p-values were corrected using Benjamini-Hochberg. Data presented as median ± interquartile range. *p < 0.05, ^a^ fecal samples were not collected when mice did not defecate voluntarily at the time of collection. Grain: grain-based diet; Syn: semi-synthetic diet.

**S3 Fig. Change in body composition in male mice over two week experimental timeline and fecal microbiota composition analyses in male mice at week 0 and week 2.** A-D) Changes in body weight and body composition of male mice on Grain, (n = 12) or semi-synthetic diet (Syn, n = 16). E. Alpha-diversity of male mice fed Grain (n = 5 – 14^a^) or Syn (n = 3 – 12^a^) diet assessed by Chao1 index. F) Beta-diversity assessed by principle coordinate analysis (PCoA), using Bray-Curtis distance metrics. G) Box plots of bacterial taxa (at genus level) at week 0 and week 2 with significant interaction, as assessed with generalized linear models with mixed effects on the sequencing counts followed by Chi Squared test. The resulting p-values were corrected using Benjamini-Hochberg. Data presented as median ± interquartile range. *p < 0.05, ^a^ fecal samples were not collected when mice did not defecate voluntarily at the time of collection Grain: grain-based diet; Syn: semi-synthetic diet.

**S4 Fig**. **Changes in body composition of male mice during 12 week experimental timeline.** Summary of body weight, body composition and dissected organ weights of male mice on Grain (n = 10) or semi-synthetic diet (Syn, n = 12) from arrival to 12 weeks. A) Changes in body weight. B) Changes in lean body mass. C) Changes in fat mass. D) Changes in percentage fat mass. E) Dissected organ weights at 12 weeks of age. Data are mean ± SEM. * p < 0.05, ^#^ 0.05 < p < 0.06. BW: body weight; BAT: brown adipose tissue; WAT: white adipose tissue; SCWAT: subcutaneous WAT; ViscWAT: visceral WAT; TG: triglycerides; BCA: bicinchoninic acid; Grain: grain-based diet; Syn: semi-synthetic diet.

**S5 Fig. Fecal microbiota composition analyses in female mice at week 0 and week 12.** A-O) Boxplots of bacterial taxa (at genus level) of female mice fed Grain (n = 8 – 12^a^) or Syn (n = 7 – 12^a^) diet with significant interactions, as assessed with generalized linear models with mixed effects on the sequencing counts followed by Chi Squared test. The resulting p-values were corrected using Benjamini-Hochberg. Data presented as median ± interquartile range. * p < 0.05. ^a^ fecal samples were not collected when mice did not defecate voluntarily at the time of collection. Grain: grain-based diet; Syn: semi-synthetic diet.

**S6 Fig. Fecal microbiota composition analyses in male mice at week 0 and week 12.** A) Alpha-diversity of male mice fed Grain (n = 7 – 13^a^) or Syn (n = 8 – 10^a^) diet assessed by Chao1 index. B) beta-diversity assessed by principle coordinate analysis (PCoA), using Bray-Curtis distance metrics. C) Boxplots of bacterial taxa (at genus level) at week 0 and week 12 with significant interaction, as assessed with generalized linear models with mixed effects on the sequencing counts followed by Chi Squared test. The resulting p-values were corrected using Benjamini-Hochberg. Data presented as median ± interquartile range. * p < 0.05, ^a^ fecal samples were not collected when mice did not defecate voluntarily at the time of collection. Grain: grain-based diet; Syn: semi-synthetic diet.

**S1 Table. Body weight and body composition of animals per batch at arrival and after two weeks.** A) for female mice. B) for male mice. Data are mean ± SEM. Differences between batches are considered significant when p < 0.05. ^a^ significantly different compared to batch 1; ^b^ significantly different compared to batch 2. Grain: grain-based diet; Syn: semi-synthetic diet.

**S2 Table. Estimate of average daily caloric intake of mice during acclimatization and throughout the study (per four week interval).** A) for female mice. B) for male mice. Average caloric intake was calculated per animal per day based on weekly weight of food consumed per cage. Data are mean ± SEM. * p < 0.05. Grain: grain-based diet; Syn: semi-synthetic diet.

**S3 Table. Body composition of Grain and Syn-fed pregnant and non-pregnant females at mating.** Grain: grain-based diet; Syn: semi-synthetic diet.

**S4 Table. Cecal short chain fatty acid (SCFA) of male mice**. Short chain fatty acid (SCFA) measurements in cecal content of Grain (n = 8^a^) or semi-synthetic diet (Syn, n = 9^a^) male mice. Data are mean ± SEM. * p < 0.05, # 0.05 < p < 0.06. ^a^ upon dissection cecums were not collected for 2 animals in Syn and one animal in the Grain diet groups, and for SCFA one cecal sample per diet group was lost. Grain: grain-based diet; Syn: semi-synthetic diet.

**Supplementary File 1. Descriptive results about incidence of fighting in male mice social housing after breeding.**

